# Nrf2 prevents diabetic cardiomyopathy via antioxidation and normalization of glucose and lipid metabolism in the heart

**DOI:** 10.1101/2022.04.04.486954

**Authors:** Ge Yang, Qihe Zhang, Chao Dong, Guowen Hou, Jinjie Li, Lingbin Meng, Xin Jiang, Ying Xin

**Affiliations:** Key Laboratory of Pathobiology, Ministry of Education, Jilin University, Changchun 130021, China; Department of Hematology and Medical Oncology, Moffitt Cancer Center, Tampa, FL 33612, USA; Department of Radiation Oncology, The First Hospital of Jilin University, Changchun 130021, China; Jilin Provincial Key Laboratory of Radiation Oncology & Therapy, The First Hospital of Jilin University, Changchun 130021, China; NHC Key Laboratory of Radiobiology, School of Public Health, Jilin University, Changchun 130021, China

**Keywords:** Diabetic cardiomyopathy, Nrf2, Lipid metabolic disorder, Glucose metabolic disorder, Oxidative stress

## Abstract

**Objective:** Metabolic disorders and oxidative stress are the main causes of diabetic cardiomyopathy. Activation of nuclear factor erythroid 2-related factor 2 (Nrf2) exerts a powerful antioxidant effect and prevents the progression of diabetic cardiomyopathy. However, the mechanism of its cardiac protection and direct action on cardiomyocytes are not well understood.

**Methods:** In this study, cardiomyocyte-restricted Nrf2 transgenic mice (Nrf2-TG) were used to directly observe whether cardiomyocyte-specific overexpression of Nrf2 can prevent diabetic cardiomyopathy and correct glucose and lipid metabolism disorders in the heart.

**Results:** Compared to wild-type (WT) mice, Nrf2-TG mice showed resistance to diabetic cardiomyopathy in a streptozotocin (STZ)-induced type 1 diabetes mouse model. This was primarily manifested as improved echocardiography results as well as reduced myocardial fibrosis, cardiac inflammation, and oxidative stress.

**Conclusion:** These results showed that Nrf2 can directly act on cardiomyocytes to play a cardioprotective role. Mechanistically, the cardioprotective effects of Nrf2 depend on its antioxidation activity, partially through improving glucose and lipid metabolism by targeting the metabolic pathways of Akt/GSK-3 β/HK-II and AMPK/Sirt1/PGC-1α.

## Introduction

In recent years, diabetes mellitus (DM) has become a serious health problem worldwide and is closely associated with the incidence of cardiovascular disease and heart failure [1, 2]. Diabetic cardiomyopathy (DCM) is currently recognized as one of the most common and serious complications of diabetes, affecting more than one-third of patients worldwide with diabetes, even if they previously did not have coronary artery disease, hypertension, or valvular heart disease [3]. The pathological changes in DCM are mainly characterized by myocardial hypertrophy and fibrosis, cardiometabolic disorders, and systolic dysfunction [4]. The occurrence and progression of DCM result from many underlying mechanisms, out of which, insulin resistance, hyperglycemia, lipotoxicity, and oxidative stress are the most crucial [5-7]. Furthermore, effective strategies to prevent or treat DCM remain lacking.

In a normal adult heart, 60%-90% of the energy demand is fulfilled by fatty acid oxidation, and the rest comes from glycolysis and lactate oxidation [8]. A normal energy metabolism of the heart is essential for its functional maintenance. However, owing to impaired insulin signaling in DCM, cardiomyocytes have a reduced ability to utilize glucose. This aggravates hyperglycemia, glucotoxicity, and accumulation of advanced glycation end products (AGEs), leading to myocardial fibrosis and cardiac dysfunction [5]. Additionally, impairment of glucose metabolism causes increased fatty acid oxidation, which in turn promotes oxidative stress and mitochondrial dysfunction [9, 10]. Subsequently, this causes an imbalance between fatty acid intake and oxidation, cardiac lipid deposition and lipotoxicity, cardiac dysfunction, and even heart failure [11]. Multiple molecular pathways participate in glucose and lipid metabolism in cardiomyocytes. For example, PPARα/PGC-1α, AMPK/Sirt1, and PPARδ/CPT1 pathways contribute to the fatty acid oxidation in cardiomyocytes [12-14], while AKT/GSK-3β, HKII/mTORC1, GLUT1, and GLUT4 signals promote glucose uptake and oxidation in cardiomyocytes [12, 15, 16]. However, these signaling pathways are impaired in the diabetic heart, aggravating cardiac glycolipid toxicity and oxidative stress, leading to cardiac dysfunction [5]. Therefore, it is essential to improve the cardiac energy metabolism to prevent DCM.

Nuclear factor erythroid 2-related factor 2 (Nrf2) is a cytoprotective transcription factor that upregulates many antioxidant genes and cytoprotective enzymes to resist oxidative stress [17]. In the late stages of DCM, cardiac dysfunction is usually accompanied by a significant decrease in Nrf2 expression. Activation of Nrf2 plays an important role in protecting against DCM [18] and heart failure by inhibiting oxidative stress [19]. Some natural molecules in green vegetables and plants, such as sulforaphane, thymoquinone, luteolin, and resveratrol, act as Nrf2 activators to protect against DCM [18, 20-22]. In addition, improving the expression of LAZ3 in diabetic mice by LAZ3 adenovirus injection can endogenously activate Nrf2 and inhibit the progression of DCM [23]. However, studies showing systemic Nrf2 activation have not clearly demonstrated whether Nrf2 exerts its cardioprotective effects by acting on cardiomyocytes or by regulating the endocrine system of the whole body in DCM. Fortunately, cardiomyocyte-restricted Nrf2 transgenic mice (Nrf2-TG) [24] can help us discover this mechanism. In addition to playing an antioxidant role by activating its downstream antioxidant genes, Nrf2 can activate AMPK and AKT signaling [25, 26]. Therefore, this study used Nrf2-TG mice to clarify whether cardiomyocyte-specific activation of Nrf2 can effectively protect against DCM, and further investigate whether Nrf2 plays a regulatory role in disorders of cardiac glucose and lipid metabolism in DM through activating AMPK and AKT signaling.

## Materials and methods

### Animals

Nrf2-TG mice were generated on an FVB background, as described previously [27]. Eight-week-old male FVB mice were purchased from Jackson Laboratory. All mice were housed in a vivarium at the University of Louisville Research Resources Center with humidified airflow and a 12/12 h light/dark cycle at a constant temperature of 22 °C with free access to rodent chow and tap water. Male Nrf2-TG and FVB wild-type (FVB WT) mice as controls were equally divided into two groups (n = 10 per group). The DM group mice were treated with five consecutive intraperitoneal (ip) injections of streptozotocin (STZ) (Sigma-Aldrich, St. Louis, MO, 40 mg/kg, dissolved in sodium citrate buffer, pH 4.5) and were named the WT/DM and Nrf2-TG/DM groups. The control groups were injected with the same volume of sodium citrate (WT and Nrf2-TG groups). Blood glucose level was measured in tail vein blood using a SureStep complete blood glucose monitor (LifeScan, Milpitas, CA), five days after the last STZ injection. STZ-injected mice with blood glucose level > 12 mmol/L were considered diabetic. Three months (3 M) and six months (6 M) after the onset of diabetes, cardiac function was examined using echocardiography. In addition, five mice in each group were euthanized for heart collection for protein, mRNA, and histopathological analyses. All procedures were approved by the Institutional Animal Care and Use Committee of the University of Louisville, which is certified by the American Association for Accreditation of Laboratory Animal Care.

### Measurement of non-invasive cardiac function

Cardiac function was measured using a high-resolution imaging system (Vevo 770, Visual Sonics, Canada) equipped with a high-frequency ultrasound probe (RMV-707B), as previously described [28]. Briefly, the mice were anesthetized with avertin (240 mg/kg, ip) and placed on a heating pad to maintain body temperature between 36–37 °C. All measurements were performed by the same operator. Echo analysis included indices of left ventricle (LV) diameters in systole (LVID; s), interventricular septal thickness in diastole (IVS; d), systolic function by ejection fraction (EF, %), and fractional shortening (FS, %).

### Serum biochemical analysis

Blood glucose was measured in tail vein blood using the Roche Accu-Chek Active blood glucose monitor. Serum triglycerides were measured before the mice were sacrificed, using an enzymatic colorimetric method (Kit No A11A01933) from Horiba ABX, France.

### Western blot analysis

Mouse heart tissue was homogenized in RIPA lysis buffer (Santa Cruz Biotechnology, Santa Cruz, CA, USA). Total or nuclear protein was extracted using a nuclei isolation kit (NUC201, Sigma-Aldrich). Protein samples were electrophoresed on a 10% sodium dodecyl sulfate-polyacrylamide gel and transferred to polyvinylidene fluoride (PVDF) membrane. The following primary antibodies from Santa Cruz Biotechnology were used: β-actin, CTGF, TGF-β1, TNF-α, Nrf2, NQO1, CAT, HO-1, and PGC-1α. Other primary antibodies used included those against p-Akt (Ser473), p-Akt2 (Ser474), p-GSK-3β (Ser9), p-LKB1 (Ser428), p-AMPK (Thr172), Akt, Akt2, GSK-3β, LKB1, AMPK, HK-II, sestrin2, Sirt1 (Cell Signaling Technology, Danvers, MA, USA), 3-NT (Millipore, Billerica, MA, USA), 4-HNE (Alpha Diagnostic International, San Antonio, TX, USA), and PAI-1 (BD Biosciences, San Jose, CA, USA). The following day, the membranes were incubated with a horseradish peroxidase-conjugated secondary antibody (Santa Cruz Biotechnology). Protein bands were visualized using an enhanced chemiluminescence detection kit (ECL, Thermo Scientific, Waltham, MA, USA). Quantitative densitometry was performed on the identified bands using the Image Quant 5.2 software (Molecular Dynamics, Inc., Sunnyvale, CA, USA).

### Cardiac histopathological analysis

To evaluate cardiac fibrosis, sections of heart tissue were stained with Sirius red, and collagen content was analyzed using a Nikon Eclipse E600 microscope, as described previously [29]. The expression of 3-NT (Millipore, Billerica, MA, USA) and 4-HNE (Alpha Diagnostic International, San Antonio, TX, USA) was determined via immunohistochemical (IHC) staining and the result was presented as positive cells/1,000 myocardial cells.

Cardiac lipid accumulation was evaluated using Oil Red O staining. Cryosections (10 μm in thickness) of cardiac tissue were incubated with Oil Red O working solution (Sigma-Aldrich) for 10 min, rinsed with 60% isopropanol, counterstained with hematoxylin (DAKO, Carpinteria, CA, USA), mounted with glycerin jelly, and observed under a light microscope (Nikon, Tokyo, Japan).

### RNA isolation and real-time PCR

Total RNA from the heart tissues was extracted using the TRIzol reagent (RNA STAT 60 Tel-Test Ambion, Austin, TX, USA). Complementary DNA (cDNA) was synthesized from total RNA using a RT-PCR kit (Promega, Madison, WI, USA) according to the manufacturer’s protocol. The primers (CAT: Mm00437229; HO-1: Mm00516005, NQO1: Mm01253561; Sestrin2: Mm00460679; β-actin: Mm00607939) were purchased from Applied Biosystems (Foster City, CA, USA). Comparative cycle time (Ct) was used to determine fold differences between samples and was normalized to an endogenous reference (β-actin).

### Biochemical measurement of lipid peroxides

Malondialdehyde (MDA) levels reflect the accumulation of lipid peroxides in heart tissue. The amount of MDA formed in the tissue was detected by the thiobarbituric acid reactivity assay, as previously described [30].

### Statistical analysis

Statistical analyses were performed using the Origin 7.5 software (Origin Lab Corporation, Northampton, MA, USA). All experimental data are presented as mean ± standard deviation (SD). Two-way analysis of variance, followed by Tukey’s test, was performed for comparison between different groups. Statistical significance was set at p < 0.05.

## Results

### Cardiomyocyte-specific overexpression of Nrf2 prevents diabetes-induced cardiac dysfunction and fibrosis

To determine whether activation of Nrf2 in cardiomyocytes protects against DCM, Nrf2-TG mice were used to establish STZ-induced type 1 diabetes model. After STZ injection, the blood glucose levels in STZ-treated mice significantly increased till 6 M. In addition, there was no difference in the blood glucose level between the WT/DM and Nrf2-TG/DM groups (Figure 1A). Compared to the WT, the levels of plasma triglycerides in WT/DM mice were significantly increased at both 3 M and 6 M time points (Figure 1B), accompanied by cardiac dilation (enlargement of LVID) and systolic dysfunction (reduced %EF and %FS), whereas the cardiac dysfunction was obviously prevented in Nrf2-TG/DM mice (Figure 1C). In addition, Sirius red staining showed increased collagen accumulation in the myocardial interstitium (Figure 1D), and increased expression of pro-fibrotic molecules CTGF and TGF-β in the WT/DM mice at 3 M and 6 M (Figure 1E). However, these diabetes-induced fibrotic responses were significantly prevented in Nrf2-TG/DM group (Figure 1D).

**Fig.1.**
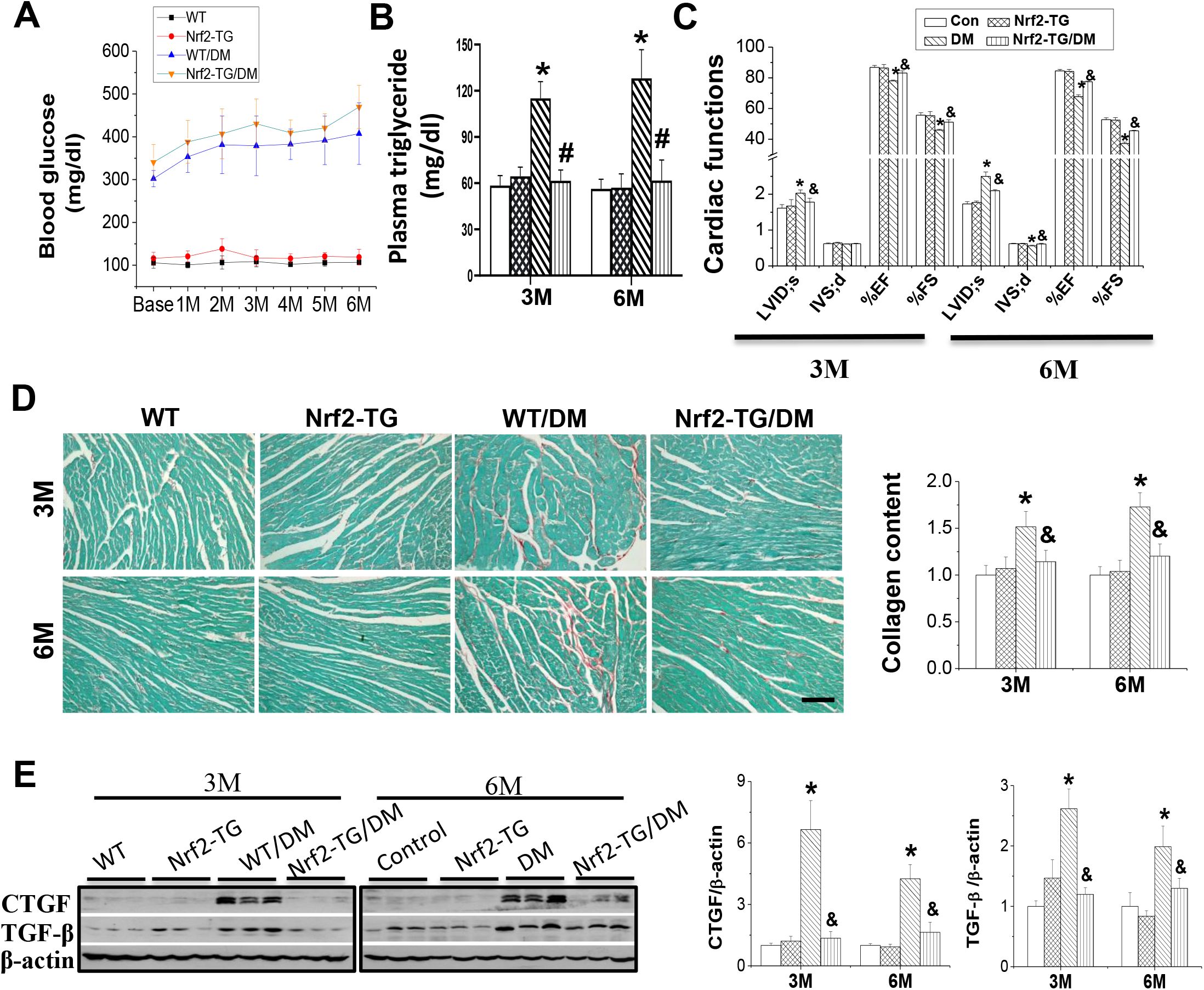
Cardiac overexpression of Nrf2 prevents metabolic disorders and myocardial fibrosis in type 1 diabetic mice. Transgenic mice with cardiomyocyte overexpression of Nrf2 (Nrf2-TG) and FVB wild type (WT) mice were injected with STZ for five consecutive days (40 mg/kg/d) and were sacrificed at 3 M and 6 M. **A**. The blood glucose level was examined with blood glucose meter. **B**. The weight was measured. **C-D**. The plasma triglyceride levels were measured by kit and Cardiac function was detected by Echo. **E**. Myocardial fibrosis was determined by Sirius red staining of collagen deposition **(bar =100μm)** and evaluating the pro-fibrotic molecules CTGF and TGF-β expression by Western blot. Data are presented as the mean ± SD (n = 5). ^*^, p < 0.05 vs. WT; ^#^, p < 0.05 vs. Nrf2-TG; ^&^, p < 0.05 vs. WT/DM.

### Cardiomyocyte-specific overexpression of Nrf2 prevents diabetes-induced cardiac inflammation and oxidative stress

Inflammation and oxidative stress play important roles in the occurrence and progression of DCM. The cardiac expression of inflammatory factors PAI-1 and TNF-α was significantly increased in the WT/DM group, but not in the Nrf2-TG/DM group, when compared with the WT group (Figure 2A). MDA content in the hearts of WT/DM mice progressively increased from 3 M to 6 M. However, overexpression of Nrf2 in the heart significantly inhibited MDA production (Figure 2B). Western blotting and IHC assay revealed that the cardiac expression of 3-NT and 4-HNE (as indices of nitrosative and oxidative damage) was significantly increased in the WT/DM group mice. On the contrary, this was almost completely prevented in the Nrf2-TG/DM group mice (Figure 2C-D).

**Fig.2.**
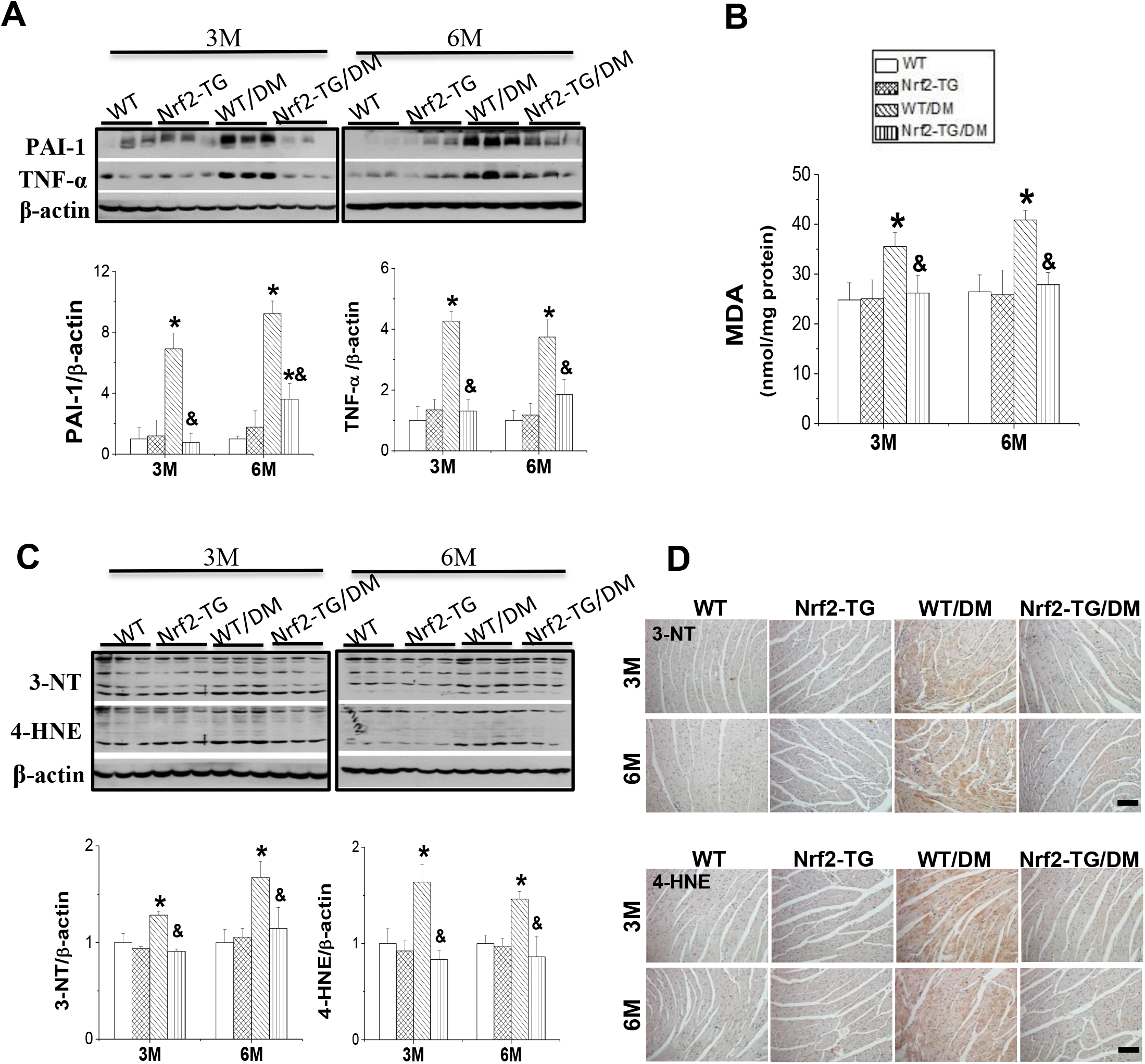
Cardiac overexpression of Nrf2 prevents cardiac inflammation and oxidative stress in type 1 diabetic mice. Mice were treated as described in **Fig.1**. **A**. The expression of inflammatory factors plasminogen activator inhibitor-1 (PAI-1) and tumor necrosis factor-α (TNF-α) were detected by Western blot. **B**. Cardiac oxidative stress was determined by MDA production. **C-D**. Western blot and immunohistochemical staining **(bar =100μm)** was used to detect the accumulation of oxidative stress markers 3 nitrotyrosine (3-NT) and 4-hydroxy-2-nonenal (4-HNE). Data are presented as the mean ± SD (n = 5). ^*^, p < 0.05 vs. WT; ^#^, p < 0.05 vs. Nrf2-TG; ^&^, p < 0.05 vs. WT/DM.

### Cardiomyocyte-specific overexpression of Nrf2 enhances the expression of cardiac antioxidant genes in mice with type 1 DM

Under both non-diabetic and diabetic conditions, Nrf2 expression in TG mice was significantly higher than that in WT mice at 3 M and 6 M (Figure 3A). Further, diabetes did not affect cardiac Nrf2 expression at 3 M, but significantly decreased its expression at 6 M (Figure 3A). However, when compared with the WT/DM group, cardiac Nrf2 expression was significantly higher in the Nrf2-TG/DM group at 3 M and at 6 M (Figure 3A). The elevated transcriptional function of Nrf2 in TG mice was also reflected in the increased expression of CAT, HO-1, and NQO1 at the mRNA and protein levels, as well as *sestrin2* at the mRNA level (Figure 3B-C). Compared to the WT group, the expression of these genes in the WT/DM group was higher at 3 M, but significantly lower at 6 M. Further, compared to the WT/DM group, the expression of these genes in the Nrf2-TG/DM group was significantly increased at 3 M and 6 M (Figure 3B-C). These results indicate that Nrf2 protects the heart from diabetes-induced oxidative damage by activating its downstream antioxidant signals.

**Fig.3.**
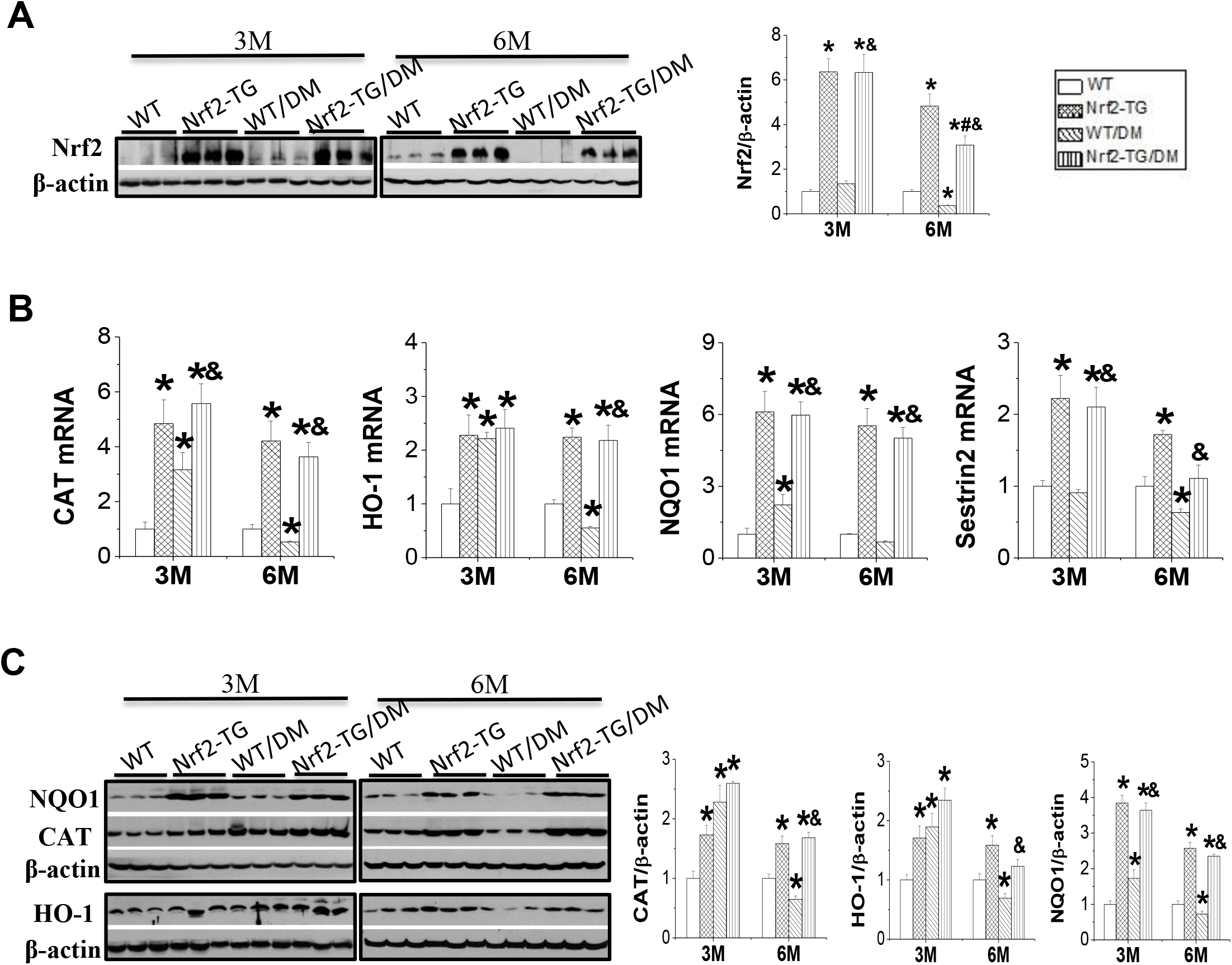
Cardiac overexpression of Nrf2 enhances the expression of antioxidant genes in type 1 diabetic mice. Mice were treated as described in **Fig.1**. **A**. Nrf2 expression was detected by Western blot. **B-C**. Nrf2 function was evaluated by detecting the expression of Nrf2 downstream genes, catalase (CAT), heme oxygenase 1 (HO-1), NAD(P)H: quinone oxidoreductase (NQO1) and sestrin2, at both mRNA and protein levels with real-time PCR and Western blot, respectively. Data are presented as the mean ± SD (n = 5). ^*^, p < 0.05 vs. WT; ^#^, p < 0.05 vs. Nrf2-TG; ^&^, p < 0.05 vs. WT/DM.

### Cardiac protection of Nrf2 occurs partly through the activation of glucose metabolism signaling via Akt/GSK-3β/HK- II

Diabetes-induced cardiac oxidative stress is associated with reduced glucose utilization due to the inactivation of Akt-mediated glucose metabolism signaling[31, 32]. Activation of the Akt/GSK-3β/HK-II signaling can improve cardiac glucose metabolism and exert cardioprotective effects [33, 34]. To explore the protective mechanism of Nrf2 on diabetic cardiomyopathy, the expression of these three glucose metabolic components was examined. Under non-diabetic conditions, the overexpression of cardiac Nrf2 did not affect the expression of HK-II and phosphorylation of Akt, Akt2, or GSK-3β (Figure 4). However, p-Akt and p-Akt2 levels were decreased in the hearts of WT/DM group at both 3 M and 6 M (Figure 4A–B). Correspondingly, GSK-3β activity increased, as shown by decreased p-GSK-3β levels (Figure 4C). In addition, HK-II expression was inhibited in diabetic mice (Figure 4D). These changes in glucose metabolic components induced by diabetes were reversed in the Nrf2-TG/DM group (Figure 4).

**Fig.4.**
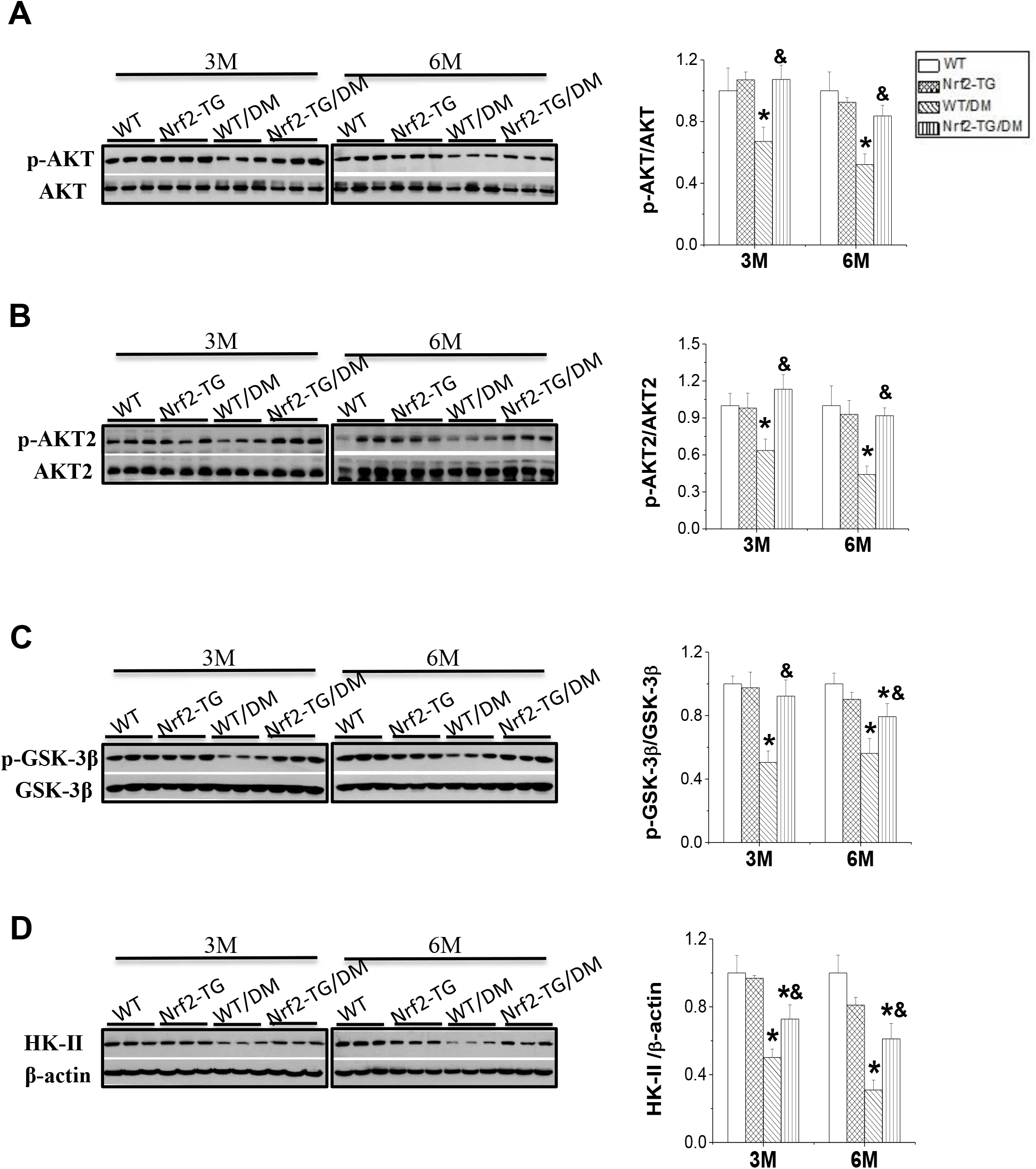
The cardioprotective effect of overexpression of Nrf2 is partly via activating AKT/GSK-3β/HK-II signaling. Mice were treated as described in **Fig.1**. **A-D**. Phosphorylation level of Akt, Akt2 and GSK-3β and expression level of hexokinase type II (HK-II) in the heart was examined by Western blot. Data are presented as the mean ± SD (n = 5). ^*^, p < 0.05 vs. WT; ^#^, p < 0.05 vs. Nrf2-TG; ^&^, p < 0.05 vs. WT/DM.

### Cardiac protection of Nrf2 occurs partly through the activation of lipid metabolism signaling via AMPK/Sirt1/PGC-1α

Cardiac lipotoxicity and oxidative stress caused by a disturbance in fatty acid metabolism are the main inducers of DCM[6, 35]. To investigate the effect of Nrf2 on cardiac lipid metabolism, cardiac lipid content was first detected by Oil Red O staining. The results showed that, compared to the WT group, cardiac lipid deposition was significantly higher in the WT/DM group, but was significantly prevented in Nrf2-TG/DM mice because of cardiomyocyte-specific overexpression of Nrf2 (Figure 5A). Nrf2 can promote the expression of sestrin2, a stress-induced metabolic regulator that activates AMPK by recruiting LKB1 to the Sesn2-AMPK complex [36, 37]. To explore how Nrf2 can regulate cardiac lipid metabolism, the activation of sestrin2 and LKB1, as well as the lipid metabolism signaling by AMPK/Sirt1/PGC-1α was studied. Overexpression of Nrf2 in the heart significantly increased the expression of sestrin2 at 3 M, but the expression level did not change at 6 M (Figure 5B). Diabetes decreased p-LKB1 expression at both 3 M and 6 M and sestrin2 expression only at 6 M (Figure 5B-C). Along with the changes in the expression levels of p-LKB1 and sestrin2, AMPK activity was inhibited (reflected by the decrease in p-AMPK) in the WT/DM group at 3 M and 6 M (Figure 5C). Correspondingly, Sirt1 and PGC-1α expression in the heart was reduced in the WT/DM group (Figure 5D). All these changes in lipid metabolic components induced by diabetes were reversed in the Nrf2-TG/DM group (Figure 5C–D).

**Fig.5.**
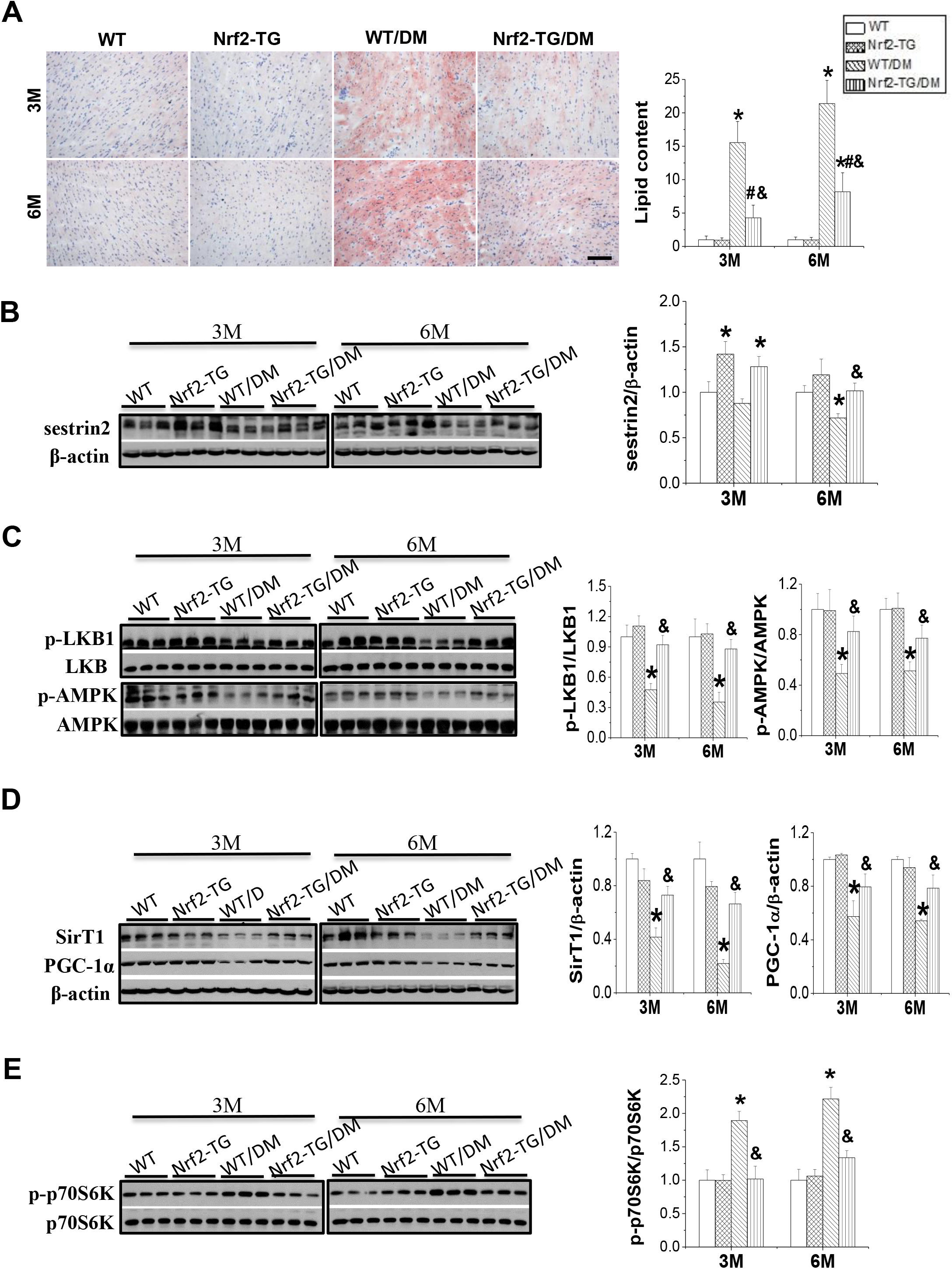
The cardioprotective effect of overexpression of Nrf2 is partly via activating AMPK/Sirt1/PGC-1α signaling. Mice were treated as described in **Fig.1**. **A**. Cardiac lipid metabolism was determined by Oil red O staining of lipid deposition. **B-E**. Cardiac sestrin2 (), Sirt1 and PGC-1α **(D)** expression level and the phosphorylation level of LKB1, AMPK **(C)** and p70S6K **(E)** was examined by Western blot. Data are presented as the mean ± SD (n = 5). ^*^, p < 0.05 vs. WT; ^#^, p < 0.05 vs. Nrf2-TG; ^&^, p < 0.05 vs. WT/DM.

### Interaction of Nrf2 with Akt and AMPK signaling

The 70-kDa S6 kinase (p70S6K) is a pivotal trigger of protein synthesis for hypertrophic change [38]. AMPK is thought to antagonize the stimulatory effect of insulin on protein synthesis by inhibiting p70S6K [9]. In addition, p70S6K inhibits Akt signaling and impairs glucose metabolism [39]. To investigate the relationship between lipid and glucose metabolism in DCM and how Nrf2 activates Akt, the phosphorylation level of p70S6K (p-p70S6K) was assessed. Activity of p70S6K was significantly higher in the WT/DM group, which could be reversed by AMPK reversal, through the overexpression of Nrf2 in the heart (Figure 5E). These results show that impaired cardiac lipid and glucose metabolism may aggravate each other in diabetes through the mediation of p70S6K, and that Nrf2 can relieve these impairments by targeting AMPK.

## Discussion

As a transcription factor that regulates antioxidant reactions, Nrf2 plays a cardioprotective role in diabetic cardiomyopathy. However, methods of systemic Nrf2 activation make it difficult to determine the specific effect of Nrf2 on cardiomyocytes [18, 40]. Therefore, we used Nrf2-TG mice, with cardiomyocyte-specific overexpression of Nrf2, to explore the direct effect of Nrf2 on cardiomyocytes in DCM and its mechanism. Our results demonstrated that specific overexpression of Nrf2 in cardiomyocytes reverses cardiac dysfunction, myocardial fibrosis, oxidative stress, inflammation, and cardiac lipid deposition caused by type 1 DM. More importantly, Nrf2 inhibits cardiac remodeling and dysfunction by activating Akt/GSK-3β/HK-II and AMPK/Sirt1/PGC-1α signals, which play important roles in regulating glucose metabolism and lipid metabolism, respectively.

Glucose uptake and metabolic dysfunction caused by insulin resistance are important pathophysiological abnormalities of DCM. In diabetic patients, insulin resistance and hyperglycemia can inhibit coronary endothelial Nitric Oxide (NO) production, increase AGE and oxidative stress in the heart, thereby promoting myocardial fibrosis, cardiac stiffness, and diastolic dysfunction [5, 41]. Akt, especially Akt2, is an important sensor of cardiac insulin signaling, which stimulates GLUT4 translocation from its intracellular location to the plasma membrane and promotes the absorption of glucose by cardiomyocytes [42]. In DCM, the expression of activated Akt is decreased, impairing the utilization of glucose by cardiomyocytes leading to cardiac dysfunction [5]. On the contrary, activation of Akt can restore heart function [43]. GSK-3β is a downstream molecule of Akt that can be phosphorylated and inactivated by Akt. In addition, activation of GSK-3β can reduce the ability of cells to synthesize and store glycogen [44] and promotes involvement in the pathogenesis of cardiac hypertrophy [45]. However, suppression of GSK-3β activity can inhibit diabetes-induced myocardial damage [46]. HK-II is a key rate-limiting enzyme in glycolysis that plays a critical role in glucose transport and phosphorylation of glucose 6-phosphate. Our previous study showed that HK-II can be negatively regulated by GSK-3β in the heart [33]. Additionally, an in vitro experiment found that inhibition of GSK-3β activity enhanced HK-II-dependent glycolysis, thereby protecting against rotenone-induced apoptosis of nerve cells [47]. Consistent with the above studies, our results also found that the activation of Akt2 (Figure 4B), caused by the overexpression of Nrf2 in cardiomyocytes, can promote the phosphorylation of GSK-3β (Figure 4C) and the expression of HK-II (Figure 4D), thereby reversing cardiac glucose metabolism disorder and cardiac dysfunction in type 1 diabetic mice. In addition, some studies have shown that the Akt/GSK-3β signaling pathway can activate Nrf2 and protect cardiomyocytes or nerve cells from oxidative stress [48, 49]. Therefore, our findings and those of others seem to support a positive feedback loop between Nrf2 and Akt2/GSK-3β in cardiomyocytes. This may further potentiate Nrf2 function in anti-oxidative stress and glucose metabolism maintenance to inhibit the progression of DCM.

Dysfunction of lipid metabolism is another major pathogenic factor in DCM. Owing to the impaired oxidation, excess fatty acids accumulate in cardiomyocytes, causing severe oxidative stress and cardiac dysfunction. Therefore, it is necessary to further explore the effects of Nrf2 on lipid metabolism. AMPK is a central regulator of energy homeostasis and can be activated by sestrin2 and LKB1 [37, 50]. Additionally, it plays an important role in regulating lipid metabolism in the mammalian heart and in resisting oxidative stress [51]. SIRT1 is a redox-sensitive enzyme of the class III histone deacetylase family. It can regulate AMPK [52] and activate PGC-1α, a transcriptional coactivator that regulates energy metabolism in an NAD(+)-dependent manner [53]. Studies have shown that activating AMPK/Sirt1/PGC-1α signaling can prevent renal lipotoxicity and inhibit mesangial cell glucotoxicity, thereby alleviating oxidative stress and apoptosis in diabetic nephropathy [54, 55]. In addition, Nrf2 can activate Sesn2 by binding to its antioxidant response element (ARE) sequence in the promoter region [36]. Based on our findings, we concluded that cardiomyocyte-specific overexpression of Nrf2 in the heart could activate AMPK by directly activating sestrin2 and LKB1, thereby preserving the activity of AMPK/Sirt1/PGC-1α signaling and inhibiting cardiac lipid deposition and oxidative stress in diabetes (Figure 5). These results are consistent with those of Dan Shao et al., who found that lipid deposition caused by impaired fatty acid oxidation is an important cause of DCM, and improving fatty acid oxidation is an effective strategy for DCM prevention [56, 57].

Under normal circumstances, the heart depends mainly on fatty acid oxidation, followed by glycolysis, for energy. However, metabolic flexibility is lost and the glucose metabolism and oxidative fatty acid pathways are both impaired in the diabetic heart [46, 58]. In addition, abnormal fatty acid metabolism in diabetes further impairs glucose metabolism, thus forming a vicious cycle and accelerating the malignant progression of DCM [57]. p70S6K, a significant downstream effector of mTOR, is involved in the development of various metabolic diseases. Several studies have shown that AMPK activation can inhibit p70S6K signaling in different tissues and cell types [59, 60], while p70S6K can inhibit Akt signaling [39]. In our study, we found that impaired lipid metabolism in the heart of patients with DCM can reduce the expression of AMPK, leading to an impaired inhibitory effect of AMPK on p70S6K, thereby probably aggravating Akt-mediated glucose metabolism (Figure 4A-B and Figure 5E). Therefore, it is considered that cardiomyocyte-specific overexpression of Nrf2 normalizes glucose and lipid metabolism by (i) directly targeting AMPK activation via its downstream genes, sestrin2 and LKB1, and (ii) indirectly reversing Akt activity via AMPK-mediated p70S6K inhibition (Figure 4A-B and Figure 5B-C, E).

In conclusion, our results suggest that overexpression of Nrf2 in cardiomyocytes preserves myocardial energy metabolism and improves cardiac function through the activation of Akt/GSK-3β/HK-II glucose metabolism signaling and AMPK/Sirt1/PGC-1α lipid metabolism signaling in DCM. Further, we show that p70S6K is the mediator of the two signaling pathways, as illustrated in Figure 6.

**Fig.6.**
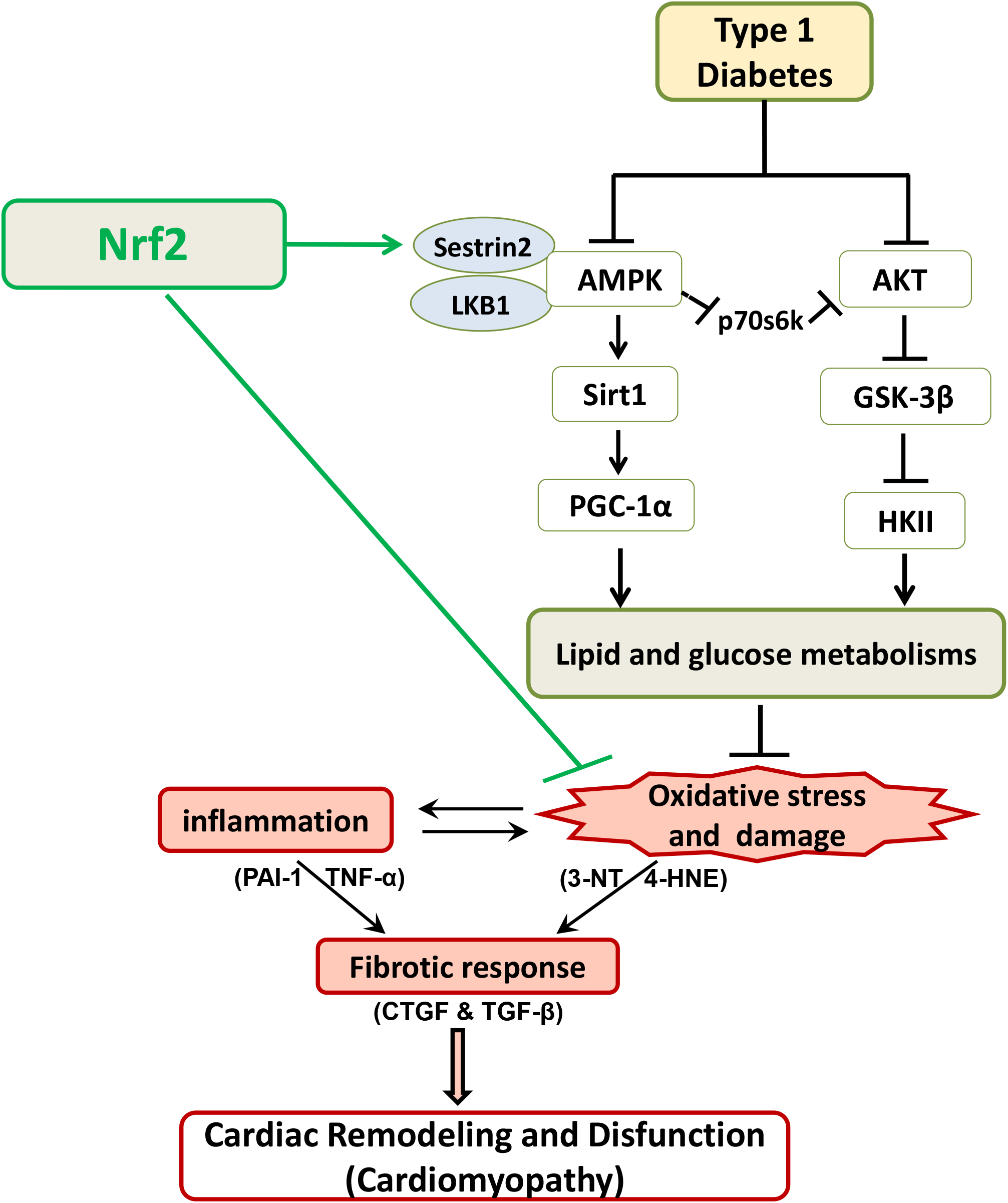
Schematic diagram of the mechanism of by which cardiac-specific overexpression of Nrf2 protects against type 1 diabetic cardiomyopathy. Type 1 diabetes causes cardiac inflammation, oxidative stress and fibrosis result in cardiac remodeling and disfunction through abnormal energy metabolism of the heart. Cardiac-specific overexpression of Nrf2 can inhibit cardiac inflammation, oxidative stress and fibrosis in type 1 diabetes through promoting normal energy metabolism via the AMPK/Sirt1/PGC-1α and AKT/GSK-3β/HK-II signaling pathway.

## Abbreviations

AGE: advanced glycation end
ARE: antioxidant response element
DCM: diabetic cardiomyopathy
DM: diabetes mellitus
EF: ejection fraction
IHC: immunohistochemistry
LV: left ventricle
MDA: malondialdehyde
STZ: streptozotocin
WT: wild-type
AMPK: AMP-activated protein kinase
AKT: protein kinase B
CTGF: connective tissue growth factor
TGF-β1: transforming growth factor 1
TNF-α: tumor necrosis factor alpha
NQO1: NAD(P)H: quinone oxidoreductase 1
CAT: catalase
HO-1: heme oxygenase 1
GLUT4: glucose transporter 4
HK-II: Hexokinase-II
PGC1-α: peroxisome proliferator-activated receptor-γ coactivator 1α
LKB1: liver kinase B1
GSK-3β: glycogen synthase kinase-3β
3-NT: 3-nitrotyrosine
4-HNE: 4-hydroxynonenal
PPARδ: peroxisome proliferator-activated receptor delta
CPT1: carnitine palmitoyltransferase 1

## Acknowledgements

We would like to thank Editage (www.editage.cn) for English language editing.

## Author Contributions

Conceptualization, X.J. and Y.X.; Methodology, G.Y., QH.Z., and C.D.; Software, GW. H., JJ.L., and G.Y.; Writing-original draft preparation, G.Y. and QH.Z.; Data Curation, G.Y. and QH.Z.; Writing-review and editing, LB.M., Y.X., and X.J.; Funding acquisition, Y.X.. All authors read and approved the manuscript.

